# Proportional modulation of proliferation and motility under 2D compressive stress depends on mesenchymal phenotype

**DOI:** 10.1101/2025.04.24.650079

**Authors:** Zacchari Ben Meriem, Moetassem Billah Meksassi, Céline Denais, Julie Guillermet-Guibert, Morgan Delarue

## Abstract

Tumor development is accompanied by strong physico-chemical modifications. Among them, compressive stress can emerge in both the epithelial and stromal compartments. Using a simple two-dimensional compression assay which consisted in placing an agarose weight on top of adherent cells, we studied the impact of compressive stress on cell proliferation and motility in different pancreatic cancer cell lines. We observed a proportional reduction of both proliferation and motility in all tested cell types, with genotypes displaying a more “mesenchymal” phenotype (high velocity-to-proliferation ratio) and others related to a more “epithelial” phenotype (low velocity-to-proliferation ratio). Moreover, “mesenchymal” cells seemed more sensitive to compression, a result that was further suggested by a TGF***β***1 induction of epithelial-to-mesenchymal transition. Finally, we measured that the change in cell proliferation was associated with a change in intracellular macromolecular crowding, which could modulate a plethora of biochemical reactions. Our results together suggest a mechanism in which all biochemical reactions related to proliferation and motility can be modulated by a change in macromolecular crowding, itself depending on the phenotype, leading to differential sensitivity to pressure.

## 1 Introduction

Tumor development is associated with genetic and physico-chemical modifications. As the tumor evolves, it accumulates genetic mutations, in particular, in pancreatic tumors cells, an activating mutation leading to hyperactive KRAS protein, such as KRAS^G12D^ mutation. This mutation is later accompanied by other mutations, such as in *TP53* encoding the p53 protein [1], which integrates, together with KRAS, a plethora of cellular processes ranging from cell proliferation to cell apoptosis. In parallel to these genomic abnormalities, solid tumors also accumulate various physico-chemical modifications. The concentration of different species such as glucose, pH and oxygen can be modified, along with the secretion of growth factors such as VEGF or TGF*β*. Similarly, tumors become more rigid with high extracellular matrix deposition, particularly in the case of pancreatic cancer [2]. Together with stromal modifications, local confined cell growth leads to the accumulation of compressive stress [3–6]. All these aspects are naturally intertwined, and collectively contribute to cancer progression.

Mechanical stresses, and in particular compressive stress, can have a myriad of effects on cell physiology, ranging from cell proliferation to migration or differentiation [7]. Notably, it has been shown in 3D that compression decreases cell proliferation and that, depending on the cell type, it can either increase or decrease cell motility [8–12]. Moreover, in a tumor setting, compressive stresses are superimposed to other cues such as chemical modifications or drug treatment, which could alter the response to compressive stress.

In this study, we investigated the effect of two-dimensional compression on cell proliferation and motility on three pancreatic cancer cell lines. We measured a proportional decrease in both cell proliferation and motility in all cell lines. However, the proportional decrease depended on the cell type, allowing us to define more “mesenchymal” cells (high velocity-to-proliferation ratio) and more “epithelial” cells (low velocity-to-proliferation ratio). We investigated the sensitivity to pressure depending on the phenotype, and explored how intracellular rheological properties such as macromolecular crowding were correlated with this sensitivity.

## 2 Results

### Two-dimensional compression of cancer cells using an agarose cushion

Adherent pancreatic cancer cells were cultured on gelatin-coated glass slides. We added a weight made of agarose directly on them to exert a compressive stress (Fig. 1A). The pressure experienced by cells was proportional to the weight and depended on both cell density and cell contact surface with the agarose [13]. We used 1% agarose to apply a pressure of *∼*100 Pa and 2% agarose to apply a pressure of *∼* 200 Pa (see Methods). As cells proliferate, the force each cell experiences decreased, and the experienced pressure decreased accordingly. However, we estimated the effect of pressure on short timescales and assumed that the pressure cells experienced was homogeneous and did not change dramatically over time - this was later reinforced by a decreased cell proliferation under pressure (Fig. 1B). Finally, the pore size of agarose was large enough such that diffusion of nutrients and growth factors was not affected by the process [14]. Notably, the imaging was performed using a holographic microscope with a large field of view (millimetric) [15], which led to images of a large number of cells simultaneously. Our method thus permitted for easy and rapid compression of adherent cells, high imaging capabilities, therefore allowing the study of the effect of compression on various physiological aspects.

**Fig. 1.**
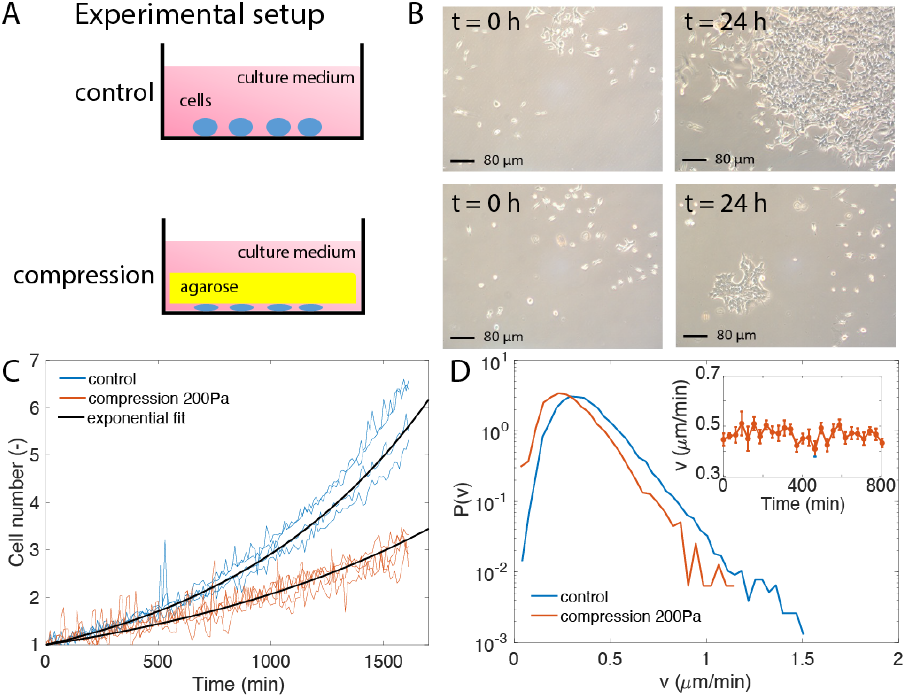
Two-dimensional compression of cells reduces both their proliferation and velocity. A. Experimental setup used to compress cells using an agarose cushion. B. Representative pictures of cells under 200 Pa and without compression. C. Normalized cell number as a function of time, and exponential fit to extract the proliferation rate. Representative curves coming from 3 different experiments. D. Histogram of single cell velocity in the field of view from 1 representative experiment. Inset: Mean cell velocity as a function of time.

### Both cell proliferation and velocity were reduced under pressure

One striking result that we observed was a concomitant reduction in cell proliferation and cell motility under compression. Indeed, for 24 hours, we can observe a significant decrease in cell number in KRAS^*G*12*D*^ mutated pancreatic cancer cells under 200 Pa (Fig. 1B). Exponential fitting of the cell number normalized to the initial cell number as a function of time, N(t)*/*N_O_ = exp(kt), led to the extraction of the proliferation rate *k* of the cell population (Fig. 1C). It appeared that the reduction in cell proliferation occurred rapidly (we can only measure a large difference within a few hours, but the decrease could be faster), and that there was no particular adaptation associated to it, as the cell number did not diverge from the exponential fit at longer timescales. In parallel, single-cell tracking enabled the measurement of instantaneous cell velocity (see Methods). We extracted the histogram of cell velocity for the different conditions and measured a decrease in mean cell velocity (Fig. 1D). To ensure that there was no temporal bias in this analysis, we measured the mean cell velocity evolution when cells were compressed as a function of time and did not observe any particular changes (Inset Fig. 1D), suggesting that the decreased cell proliferation occurred rapidly and that there was no adaptation to it.

### Proliferation decreases linearly with pressure, and depends on the genotype

We extended our experiments to three different epithelial cell lines bearing specific mutations that are present during pancreatic cancer development or alter it: KRAS^*G*12*D*^ alone or in combination with a p53^*R*172*H*^ or a PI3K*β*^*in*^ activating mutation [1]. We observed, in all cell lines, a progressive decrease in cell proliferation under 2D compression (Fig. 2A). The decrease in cell proliferation appeared to be linear with pressure. Using a linear fit, we could extract a characteristic pressure *P*_*c*_ related to the sensitivity of cell proliferation to pressure: 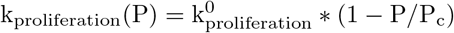. The smaller *P*_*c*_ was, the more sensitive the cells were to pressure. We noticed that *P*_*c*_ depended on genotype, and that KRAS^*G*12*D*^ mutated cells were more sensitive to pressure than in combination with other genetic alterations (Fig. 2B).

**Fig. 2.**
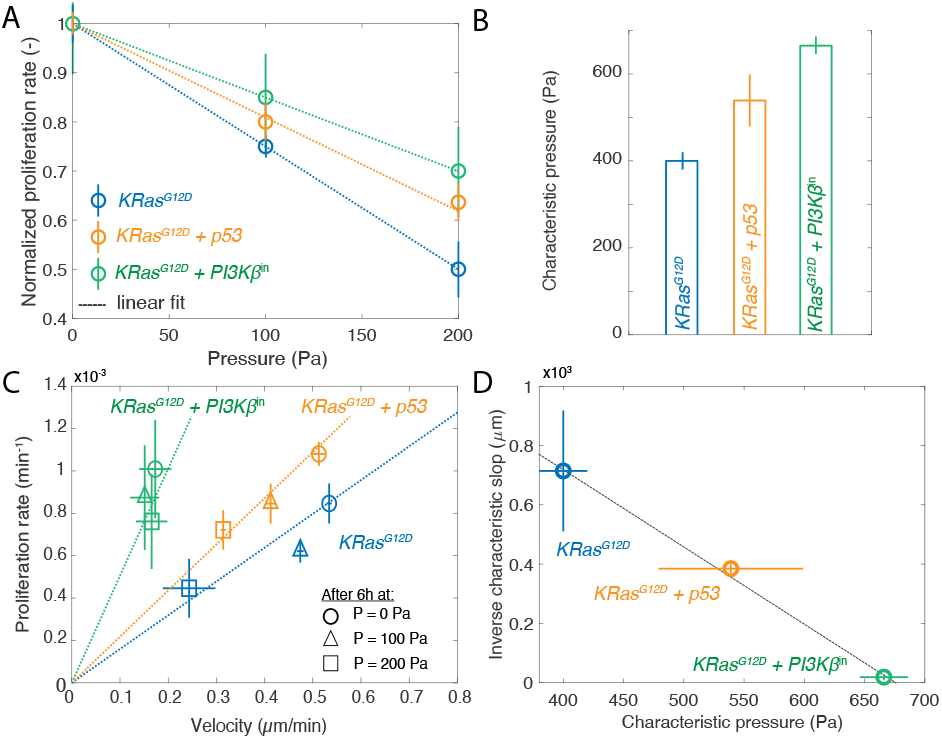
Effect of compression on proliferation and velocity in different cell lines. A. Linear reduction of proliferation rate as a function of compressive stress. B. Characteristic pressure of proliferation reduction for the different cell lines. C. Proportional reduction of proliferation and velocity as a function of pressure in different cell lines. D. Correlation between the proportional reduction of proliferation and velocity with the characteristic pressure of proliferation reduction. For each point, at least 3 independent replicates were performed. Error bars denote mean *±* standard error of the mean.

### Proportional decrease of proliferation and velocity under pressure

We equally measured the cell velocity as a function of compressive stress for the three different genotypes. Even if the data were noisier, we similarly observed an almost linear decrease in cell velocity. As a consequence, when we plotted the proliferation rate as a function of cell velocity for the different genotypes and pressure conditions, we observed a proportional decrease of proliferation rate and velocity as pressure increased (Fig. 1C). In this graph, we could observe that KRAS^*G*12*D*^ cells have higher motility relative to their proliferation rate than KRAS^*G*12*D*^ + PI3K*β*^*in*^ cells, for example. This could indicate a more pronounced “mesenchymal” phenotype, in opposition with a more “epithelial” phenotype which would correspond to larger cell proliferation and lower cell motility.

### Correlation between “mesenchymal” and “epithelial” phenotypes with pressure sensitivity

We then calculated the slope of the proportionality between the proliferation rate and the velocity (dashed line in Fig. 2C), and plotted its inverse (the velocity-to-proliferation ratio) as a function of the characteristic pressure estimated in Fig. 2B. Interestingly, we observed a strong anticorrelation between the velocity-to-proliferation ratio, which could be interpreted as the strength of the mesenchymal phenotype, and the sensitivity to pressure (Fig. 2D): more “mesenchymal” cells appeared more sensitive to compression, both in terms of proliferation and motility, than more immobile “epithelial” cells.

### Induction of a mesenchymal phenotype increases pressure sensitivity in a time-dependent manner

Even if KRAS^*G*12*D*^ cell line displayed a more “mesenchymal” phenotype, it remained an epithelial cell line. We next sought to induce an “epithelial-to-mesenchymal” transition of these cells by treating them with TGF*β*1 [16]. We investigated the effect of this chemical inducer in terms of its impact on cell proliferation and cell velocity, rather than its molecular effects. Intriguingly, we measured that TGF*β*1 induction increased both cell motility and cell proliferation on short timescales (hours). The effect seemed to be additive to the effect of pressure, as the increase in both proliferation and velocity was observed with a similar increase under pressure (Fig. 3A). Cell adaptation to TGF*β*1 occurred on longer timescales, where a clearer mesenchymal phenotype appeared, with a decrease in cell proliferation, while retaining the increase in cell velocity. When estimating the characteristic pressure related to the decrease in cell proliferation, we measured that it decreased to *P*_*c*_ *∼* 300*Pa*, consistent with our other observations that a more “mesenchymal” phenotype would be more sensitive to pressure.

**Fig. 3.**
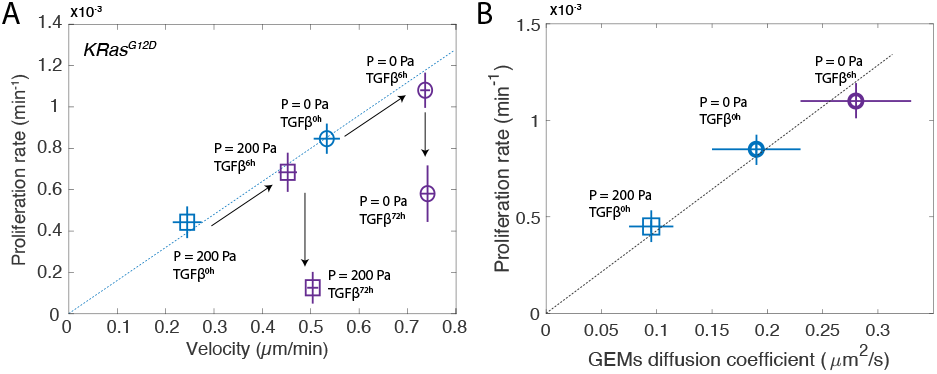
Additive effect of TGF*β*1 and compressive stress under pressure. A. After 6h, TGF*β*1 increases both cell proliferation and velocity, keeping the proportionality between proliferation and velocity. At longer timescales, proliferation decreases, while cell velocity remains unchanged. B. The change in intracellular diffusion of genetically-encoded nanoparticles correlates with the change in proliferation under pressure and TGF*β*1 induction. Error bars denote mean *±* standard error of the mean for cell proliferation, and mean *±* standard deviation for diffusion coefficient, in N *≥* 3 independent experiments.

### The linearity between proliferation and velocity is correlated with the intracellular rheological properties of the cell

Finally, we wished to investigate the linear relationship between proliferation rate and cell velocity. We reasoned that, being conserved among several cell lines and changing proportionally for both a mechanical stress and a short pulse of TGF*β*1 induction, this linearity could be related to intrinsic cellular parameters. We thus decided to measure how the intracellular macromolecular crowding, a key cellular parameter that is known to impact a plethora of cell processes [17, 18] from biogenesis [19, 20] to the polymerization of microtubules [21]. We genetically encoded tracer nanoparticles called GEMs [22] and measured their diffusion as a function of mechanical compression and TGF*β* induction. We observed a striking linear relationship between proliferation rate and GEMs diffusion (Fig. 3B), suggesting that cytoplasm fluidity was indeed related to the observed phenotypes (proliferation and velocity) observed in our experiments.

## 3 Conclusion and Discussion

To investigate the effect of 2D compression on pancreatic cancer cells, we developed a simple compression assay consisting in applying a constant agarose weight on top of adherent cells. We measured that compressed cells proliferated and moved more slowly with increasing compression, and that the sensitivity to compression seemed to be dependent on the “epithelial / mesenchymal” state of the cell, as, even in these epithelial cells, a more “mesenchymal” phenotype was associated with a larger sensitivity to pressure. Interestingly, we observed an increase in macromolecular crowding under compression, which could explain the reduction of proliferation and motility under pressure.

Even if the effect of compressive stress on cell proliferation in 2D has been much less explored than in 3D, this result is not surprising, as it has been repeatedly observed experimentally. However, it is interesting to note that, depending on the genotype, cells are more or less sensitive to compression. This characteristic needs to be placed back into a tumor setting, investigating how such mutations could affect tumor progression in the context of mechanical stress. In particular, recent results suggest that the status of PI3K activity could be correlated to the responsiveness of cells to compressive stress [13].

The effect of compression on velocity is more controversial, with some studies showing an increase of motility, while others show a decrease [8–12]. This discrepancy could originate not only from the cells used in the studies but also from the experimental conditions. For instance, some studies investigating the effect of compression on motility alone tend to study them in PBS to dramatically decrease cell proliferation [8,9]. However, doing so could imbalance resource allocations in cells and lead to different results. In other studies, this is not compression that is controlled but a 2D confinement [12]: it is thus impossible to estimate the level of stress in these experiments without knowing the cellular mechanical properties, such that cell lines displaying increased velocity in this setting could display a lower one in ours.

A striking result of our study is the proportionality between the decrease in cell proliferation and cell velocity. Although the strength of this decrease depended on the cell line, it nonetheless existed. We reasoned that such a proportional decrease could stem from a global regulation of biochemical reactions. If all reaction rates decrease/increase, we should observe a corresponding modulation of both proliferation and motility. Macromolecular crowding, which relates to the intracellular rheological properties of the cells, can modulate all biochemical reactions by affecting the affinity between substrates as well as their diffusivity [17, 18]. We observed that crowding increased under compression and decreased under a short-term TGF*β*1 induction, such that we measured a strong correlation between proliferation rate and intracellular diffusivity of genetically-encoded tracer nanoparticles (GEMs) [22]. This result suggested that pressure could modulate crowding, which, in turn, can modulate fundamental processes such as proliferation and motility. Notably, this proportionality was lost after the cells had adapted to TGF*β*1. However, in this case, the cell phenotype has changed and the proteome and resource allocation are likely modified.

To conclude, our results together suggest a mechanism in which compressive stress increases crowding, which could decrease biochemical reactions in cells. This modulation would depend on the cell genotype, with some cell lines being more responsive than others. Interestingly, less responsive cells have been associated with an increased autophagy: this process could de-crowd the cells, explaining the differential sensitivity [13]. This part would be interesting to investigate in a future study. Finally, it is interesting to observe that cells that could be defined as more “mesenchymal” seemed to be more responsive to compression than cells that could be defined as more “epithelial”. While compression can lead to differentiation of mesenchymal cells and promote the production of collagen matrix, participating in bone regeneration and healing [23], it remains to be investigated if this differential sensitivity of mesenchymal cells to compression could have implications during tumor development [16].

## 4 Material and Methods

### Cell provenance and cell culture

All of the pancreatic cancer cell lines were collected from murine pancreatic tumors bearing specific mutations. The A338 pancreatic cancer cell line displays an activating KRAS oncogene mutation (KRAS^*G*12*D*^), the B385 cell line bears the KRAS^*G*12*D*^ mutation as well as a PI3K*β* inactivating genetic alteration, and the R211 cell line encompasses the KRAS^*G*12*D*^ mutation together with the p53^*R*172*H*^ gain of function mutation [1, 24]. These pancreatic cancer cell lines were cultured in DMEM Glutamax, pyruvate (Gibco, ThermoFisher Scientific) supplemented with 10% Fetal Bovine Serum (Sigma-Aldrich) and 1% penicillin-streptomycin (Sigma-Aldrich) at 37^*°*^C with 5% CO_2_.

To create all GEM cell lines, the construct pUBC-Pfv-Sapphire-PURO, which was a gift from Liam Holt (Addgene plasmid number 203651), was obtained from Addgene. DH5*α* competent bacteria containing the plasmid of interest were expanded on 10*µ*g/mL ampicillin-supplemented Luria Broth plates, collected, and amplified in LB/ampicillin liquid subcultures. The GEMs encoding plasmid was subsequently purified using the GeneJET Plasmid miniprep kit (Thermo Scientific).

6-well plates were coated with 0.1% of gelatin (Stemcell Technologies), and 3×10^5^ cells were plated into each well. Upon reaching 70-80% confluency, cells were transiently transfected with 2*µ*g of the construct coding for GEMs nanoparticles using jetPRIME transfection reagent (Polyplus) according to the manufacturer’s instructions.

The A338 GEMs cell line was selected with 1*µ*g/mL of puromycin (Gibco) and then maintained in culture using 0.3*µ*g/mL of puromycin antibiotic. These concentrations used for selection were determined by antibiotic selection kill-curve. Cells were then cultured on *µ*-Dish 35mm imaging dishes with glass bottoms (Ibidi) overnight. Following attachment, cells were treated with TGF*β* (recombinant Human TGF-1 10µg (HEK293 derived) ref100-21-10UG (Preprotech), used at a concentration of 5ng/mL) reconstituted in citric acid 10mM and resuspended with 1X phosphate-buffered saline (PBS) and 0.1% bovine serum albumin (BSA).

### Compression system

Cells cultured in 2D were subjected to mechanical compression by placing a 1-2% agarose pad directly on top of the cell layer, applying a final compressive stress of 100-200 Pa [13]. The control was simply the same cells without any weight applied. To prepare the agarose pads, UltraPure Agarose 1000 (Invitrogen) was dissolved in 1X phosphate-buffered saline (PBS) and heated in a microwave to ensure complete dissolution. The molten agarose (40 mL of 2% solution) was then poured into a Petri dish and allowed to solidify. Round agarose pads were subsequently cut out using the cap of a 50 mL Falcon tube.

### Image acquisition and analysis

Cells were imaged every 10 minutes using a holographic microscope Cytonote 6W (Iprasense). Single-cell segmentation and trajectories were performed and extracted using Image J pluggin Trackmate (LoG detector setting with estimated object diameter 14 pixel, quality threshold 1 with sub-pixel localization). Mean velocities were determined from the trajectories throughout the experiment.

Control, compressed, and TGF*β* (5ng/mL) treated cell samples were cultured on *µ*-dishes with glass bottom allowing high-resolution acquisitions. Imaging was performed using a Leica DM IRB microscope, equipped with a Yokogawa CSU-X1 spinning-disk confocal and a Hamamatsu ORCA-Flash4.0 v3 sCMOS camera, with a 63x objective. GEMs nanoparticle movements were captured using a 488 nm laser at full power, with 200 images continuously acquired over 2s at a frame rate of 100 Hz.

Particle tracking and trajectory analysis were conducted in FIJI using the MOSAIC plugin, which enabled the extraction of individual particle trajectories. Time-averaged mean square displacement (MSD) was calculated for each trajectory, with the first 10 points (100 ms) fitted to a linear model to obtain single-particle diffusion coefficients. Mean values and standard errors were then computed from the extensive data collected, following the methodology of Ref. [22].

## 5 Acknowledgement

This work was supported by Inserm Plan Cancer PCSI (Press-Diag-Therapy and MechaEvo projects) as well as ERC Starting grant Under-Pressure grant agreement number 101039998). Views and opinions expressed are, however, those of the author only and do not necessarily reflect those of the European Union or the European Research Council. Neither the European Union nor the granting authority can be held responsible for them.

## 6 Author contribution

ZBM, JGG and MD conceived the experiments. ZBM and MD realized, and analyzed the compression experiments. CD created the GEM cell line. MBM realized and analyzed the GEMs experiments. MD wrote the manuscript. All authors proofread the manuscript.

## 7 Data availability

Data will be deposited on Zenodo repository after publication.

## 8 Literature

